# A Simplified Function-First Method for the Discovery and Optimization of Bispecific Immune Engaging Antibodies

**DOI:** 10.1101/2022.08.17.504342

**Authors:** Alex Shepherd, Bigitha Bennychen, Anne Marcil, Darin Bloemberg, Rob Pon, Risini Weeratna, Scott McComb

**Affiliations:** Human Health Therapeutics Research Centre, National Research Council, Canada; Centre for Infection, Immunity and Inflammation, University of Ottawa, Ottawa, Canada; Department of Biochemistry, Microbiology, and Immunology, University of Ottawa, Ottawa, Canada

## Abstract

Bi-specific T-cell engager antibodies (BITEs) are synthetic soluble molecules derived from antibodies that induce active contact between T-cells and other target cells in the body. BITE therapeutics have shown great promise for the treatment of various forms of cancer; however, the current development process for BITEs is time consuming and costly. BITE development requires empirical testing and characterization of the individual antigen binding domains, followed by extensive engineering and optimization in bi-specific molecular format to generate a molecule with strong biological activity and appropriate characteristics for clinical development. Here, we sought to create a cost efficient high-throughput method for creating and evaluating BITEs using a simplified function first approach to identify bioactive molecules without purification. Using a plasmid with a modular structure to allow high efficiency exchange of either binder arm, we established a simple method to combine many novel tumour-targeting single chain variable (scFv) domains with the well-characterized OKT3 scFv CD3-targeting domain. After generating these novel plasmids, we demonstrate two systems for high throughput functional screening of BITE molecules based on Jurkat T cells (referred to as BITE-J). Using BITE-J we evaluate four EGFRvIII BITEs, identifying two constructs with superior activity. We then confirmed this activity in primary T cells, where novel EGFRvIII-BITEs induced T cell activation and antigen selective tumor killing. We also demonstrate that we can similarly exchange the CD3-interacting element of our bi-modular plasmid. By testing several novel CD3-targeting scFv elements for activity in EGFRvIII-targeted BITEs, we were able to identify highly active BITE molecules with desirable properties for downstream development. In summary, BITE-J presents a low cost, high-throughput method for the rapid assessment of novel BITE molecules without the need for purification and quantification.

## Introduction

Monoclonal antibody (mAb) technology can be used to create biological molecules with high affinity and specificity for antigenic targets on the surface of cells. Through binding, such antibodies can induce a variety of direct and indirect biological effects, such as agonistic or antagonistic receptor modulation^1^. In addition to these direct effects, antibodies can simultaneously recruit immune cells through interactions with the antibody Fc domain, creating a connection between target cells and immune cells that can lead to phagocytosis or antibody-dependent cytotoxicity^1^. Synthetic immunology approaches have been developed to broaden the effects of antibodies and derivative molecules in immune cell activation, including the development of bi-specific T-cell engager (BITE) antibodies that can induce strong antigen-targeted T cell responses^2^. The most clinically advanced of such therapies is Blinatumomab, a CD19xCD3 antibody^3^ used in the treatment of acute lymphoblastic B-cell leukemia^4^.

To further broaden the types of cancer that can be effectively targeted using BITE therapeutics, optimize their biological activity, and explore additional modalities for bi-specific antibodies, there is a need for cost-effective and efficient high-throughput platforms for discovering/developing novel BITE molecules. BITE development begins with the identification of two antigen-binding molecules, most typically single chain variable fragments (scFv). One scFv must bind an antigen on a target cell, with the other binding to an immune cell, most commonly recognizing CD3 on T cells.^5^ These scFvs are connected using a linker, commonly composed of a flexible amino acid sequence (GGGGS) length as seen in Blinatumomab^6^, which can also be repeated to create a longer flexible linker domain (GGGGS x 2, 3 etc..). In the presence of a BITE molecule, T cells and target cells expressing the target antigen should form a strong interaction, prompting both T cell activation and target cell killing^7,8^.

While assessment of common antibody quality attributes such as avidity and monovalent affinity are critical to understand the immunochemical properties of a mAb, it is not well understood what antibody properties are most desirable for development of a bi-specific immune cell recruiting antibody. Other methods such as dock and lock (DNL) have been devised to create multi-specific antibody complexes by fusing antibodies through anchoring proteins^9^, methods such as these are complex, require purified antibody proteins, and create molecules of large size, potentially complicating downstream process development and limiting their potential applications. Here we outline a complete, high-throughput and easy to use method for generating novel bi-specific antibody plasmids, producing BITE molecules and performing activity screening in human Jurkat T cell line that does not require antibody preparation, purification and characterization. This allows for immediate functional BITEscreening of BITE candidates following DNA synthesis or assembly, reducing the time and cost associated with finding a lead candidate.. Using the tools and techniques outlined in this paper, researchers can quickly discover and optimize ideal candidates for further clinical development or incorporation into more complex synthetic therapeutics.

## Materials and Methods

### Construction of pBITE modular plasmid

The plasmid design is modeled on blinatumomab^10^ and contains a IgG1 signal peptide, modularized CD19-scFv (OKT3HD37), linker, modularized CD3-scFv (OKT3), and P2A-NeonGreen (see Supplemental Sequence document for DNA and amino acid sequences used). To build the pBITE plasmid, CAR-specific domains were removed from pSLCAR-CD19 (Addgene #135992), leaving a linearized plasmid containing a short EF1a promoter and a P2A-NeonGreen reporter sequence. Next, a custom DNA fragment was designed and synthesized (Twist Bioscience, USA) coding for the HD37 anti-CD19 scFV, linker, OKT3 anti-CD3 scFv and adapter sequences to allow cloning into the linearized plasmid using Gibson assembly^11^. Gibson assembly was performed using either the GeneArt™ Gibson Assembly HiFi Master Mix (ThermoFisher A46627) or an in house “DIY Gibson” assembly mixture using the RFC57 recipe (^12^. The final plasmid incorporates modular restriction sites to allow swapping of the tumour engager arm (using BpiI) and the CD3 binding arm (using Esp3I), as well as the linker arm between the two. The resultant plasmid was transformed into DH5a chemically competent *E. coli*. To confirm successful cloning, individual transformed colonies were analyzed with PCR using backbone or scFv-specific primers. Colonies which contain predicted PCR products are grown overnight in normal lysogeny broth (LB) with 1% ampicillin. Plasmid DNA was purified using standard mini-plasmid or midi-plasmid purification kits and then sequenced via Sanger sequencing to ensure accurate sequence construction.

### Tumor antigen binding domain or CD3-binding domain exchange

Replacement scFv sequences are typically between 400-600 bp long. When designing them for “Golden-Gate” simultaneous restriction/ligation, the 5’ and 3’ ends are structured depending on the cloning site used. For swapping of the N-terminal tumour-antigen binding domain using BpiI the following sequences were used for cloning: 5’ –NNN**GAAGAC**NNAGGA - *VH or VL sequence – Linker – VH or VL sequence –* NNCCTTNN**GTCTTC**NNN-3’. For swapping of the C-terminal immune cell engaging domain using Esp3I 5’ –NNN**CGTCTC**NAAGT - *VH or VL sequence – Linker – VH or VL sequence –* GCTAN**GAGACG**NNN-3’. In this manuscript a VH-Linker-VL orientation was used for all scFvs tested.

To swap either scFv, the following reaction conditions were used: 250ng of pBITE, 50ng of new scFv, 0.25ul BpiI or Esp3I, 0.25ul T4 ligase, 4ul T4 reaction buffer and filled to 20ul with molecular grade H_2_O. This reaction is run on a thermocycler under the following protocol: (37°C for 10 min, 16°C for 10 min) x cycles, 37°C for 60 min, 80°C for 5 min, and 4°C indefinitely. 5ul of this reaction was heat shocked into DH5α chemically competent *E. coli* and then plated on ampicillin containing agar plates. To confirm successful cloning, individual transformed colonies were analyzed with PCR using backbone or scFv specific primers. Colonies which contained predicted PCR products are grown overnight in normal lysogeny broth (LB) with 1% ampicillin. Plasmid DNA was purified using standard mini- or midi-prep purification kits (Qiagen, USA). It is recommended that purified plasmids are sequenced to ensure correct construction. Sequencing primers used for Sanger sequencing of the plasmid were as follows: EF1a-promoter-Seq-F: CGCAACGGGTTTGCCGCCAGAACACAG, BM-seq-link-F: CCTTGGAGGAGGCGGAAGTC, NeonG-seq-R: TCGAAGTCCACGCCGTTGATG.

EGFRvIII-specific scFv sequences used for cloning can be found in ^13^ and ^14^ (F263-4E11, F265-5B7, F269-3D12, and F271-1D2).

### Cell lines

All cell lines were monitored regularly for mycoplasma contamination using in-house PCR assay^15^. In preparation of cell assays, healthy cultures of Jurkat E6-1 cells (ATCC#TIB-152) were maintained in complete RPMI (RPMI1640 supplemented with 10% FBS, 2mM L-glutamine, 1mM sodium pyruvate and 100 µg/mL penicillin/streptomycin) with cell density between 0.25 and 1×10^6^ cells/mL for several weeks; we have found this to be critical for consistent results in Jurkat activation assays. All target cell lines described in this paper were modified using Nuclight-Red Lentiviral reagent (Cat#4625, Sartorius, USA) to generate stable red-fluorescent cells which can be easily differentiated from effector cells in FACS or live microscopy analyses. Specific target lines used were as follows: Raji (CD19+, ATCC#CCL-86), Nalm6 (CD19+, DSMZ#ACC128), MCF7 (CD19-, ATCC#HTB-22), U87MG-vIII (EGFRvIII+, a gift from Prof. Cavnee, Ludwig Cancer Institute, USA), U87MG (EGFRvIII−, also a gift from Dr. Cavnee), DKMG (EGFRvIII^low^, DSMZ#ACC277). Target cell lines were grown in varying media conditions, as recommended by cell repository.

### Jurkat Electroporation protocol

In preparation of cell assays, healthy cultures of Jurkat cells with cell density between 0.25 and 1×10^6^ cells/mL were maintained in culture for several weeks. Prior to electroporation, a recovery plate was prepared using pre-warmed RPMI 1640 supplemented with 20% fetal bovine serum and 100 ug/ml of L-glutamate. Jurkat cells were pelleted via centrifugation, then resuspended in 100ul of 1SM buffer (as per Chicaybam et al^16^) and 2ug of plasmid and brought to room temperature. Cells were then placed into a 0.2cm electroporation cuvette (Cat#1652086, Bio-Rad, USA) and electroporated using a Lonza Nuceleofactor Electroporator using X4 settings. Cells were then immediately transferred to the recovery plate. Cells were allowed to recover for 4 hours prior to analysis or use in an assay.

### HEK293 transfection protocol

HEK293 cells were plated in a 6 well plate at approximately 150k cell per well and allowed to grow overnight. Cells appeared healthy and cell confluence was around 60% when transfection was performed. The following transfection mixture was prepared: 1.2ml of serum free DMEM 200ul/well, 9ug/well linearized polyethyleneimine (PEI; Cat#765090-1G, Sigma-Aldrich, USA), and 12ug (2ug/well) of pBITE plasmid. The tube containing the transfection mix was vortexed and incubated at RT for 10 minutes. 200ul of the solution was added drop wise to each well incubated at RT for 20 minutes. Cells were then incubated at 37°C for 4 hours. After 4 hours, media was removed and replaced with 2-3ml of complete DMEM media. After 5-7 days, transfected HEK293T supernatant was removed and filtered using 0.45-micron filter to remove any excess cells. BITE-containing supernatant was then frozen at −80°C and thawed prior to BITE-Jurkat assay. We confirmed that freezing resulted in no loss of BITE activity for the active BITE molecules reported in this manuscript.

### BITE-Jurkat Activation Assay

If adherent target cells were used, cells were first treated with accutase (Cat#A1110501, Thermo Fisher, USA) to create a single cell suspension. Jurkat cells and target cells were then resuspended in complete RPMI and placed in 96-well plates at varying ratios of Jurkat and target cells as shown in the text. Unless otherwise stated in the text, 50ul of BITE containing HEK supernatant were added to appropriate wells. Complete media was then added to equalize plating volume. Plates were incubated at 37°C, 5% CO_2_ for either 16 or 40 hours. Plates was then stained with anti-CD69 APC (Cat#340560, BD Bioscience, USA) at 0.25ul per well in no-wash format. Plates were then incubated in the dark at RT for 15 minutes and analyzed via flow cytometry using BC Fortessa device (BD Bioscience, USA).

### Primary Human T cell culture preparation

To prepare T cells, healthy donor blood samples were obtained under appropriate safety and ethical approval by the Ottawa Hospital Research Institute (Ottawa, Canada). Whole blood was diluted 1:1 with Hank’s balanced salt solution (HBSS) and PBMCs were isolated by Ficoll-Paque™ density gradient centrifugation, centrifuging for 20 min at 700 × g without applying a brake. The PBMC interface was carefully removed by pipetting and was washed twice with HBSS by stepwise centrifugation for 15 min at 300 × g. PBMCs were resuspended and counted by mixing 1:1 with Cellometer ViaStain™ acridine orange/propidium iodide (AOPI) staining solution and counted using a Nexcelom Cellometer Auto 2000 (Nexcelom BioScience, Lawrence, Massachusetts, USA). T cells from were then activated with Miltenyi MACS GMP T cell TransAct™ CD3/CD28 beads and seeded 1×10^6^ T cells/ml in serum-free StemCell Immunocult™-XF media (Cat#10981, StemCell Technologies, Vancouver, Canada) with 20U/ml clinical grade human IL-2 (Novartis). T cells were then polyclonally expanded for 7 to 10 days in culture before cryopreservation of aliquots using complete media + 10%DMSO.

### BITE-Induced Target Cell Killing Assessment via Live Fluorescence Microscopy

Cryopreserved polyclonally expanded primary human T cells were thawed on the day of the experiment and counted and assessed for viability using Nexcelom Cellometer Auto 2000 (Nexcelom BioScience, Lawrence, MA, USA). Human T cells and red-fluorescent target cells were then resuspended in Immunocult-XF media at appropriate concentration. Cells were then transferred to a 96-well culture plate at 10 000 T cells per well and 2000 target cells per well. Plates were then placed in an S3 Incucyte Live Cell Imagine system (Sartorius, USA) and incubated at 37°C and 5% CO_2_ for up to 140 hours. Images were taken every 2 to 4 hours. Target cell growth was tracked using automated assessment of red fluorescent area percentage using the accompanying Incucyte analysis software. Final data was assembled and analyzed using Graphpad Prism.

## Results

### Creating a modularized Bispecific plasmid

Previously we reported on a Jurkat based platform for screening novel chimeric antigen receptor molecules^17^, here we wished to create a similarly flexible plasmid which could be used to screen single-chain variable fragments (scFv) for activity in a soluble bi-specific antibody molecules (Figure 1a). We based our plasmid design on the amino acid sequence for blinatumomab^10^, adding type-IIs restriction sites to allow for simplified cloning of either binding arm of the molecule (Figure 1b). The modularized blinatumomab biosimilar plasmid (pBITE) allows either end of the BITE to be swapped for an alternative scFv in a single golden gate reaction (Figure 1b), as well as exchange of the linker connecting both scFv domains if desired. This system can be used to easily customize the BITE design, allowing for efficient construction of a variety of BITEs and BITE compositions for testing. Internal testing shows highly efficient exchange of both target antigen scFv and T cell antigen scFv can be achieved consistently using single pot restriction ligation as described in the methods section.

**Figure 1:**
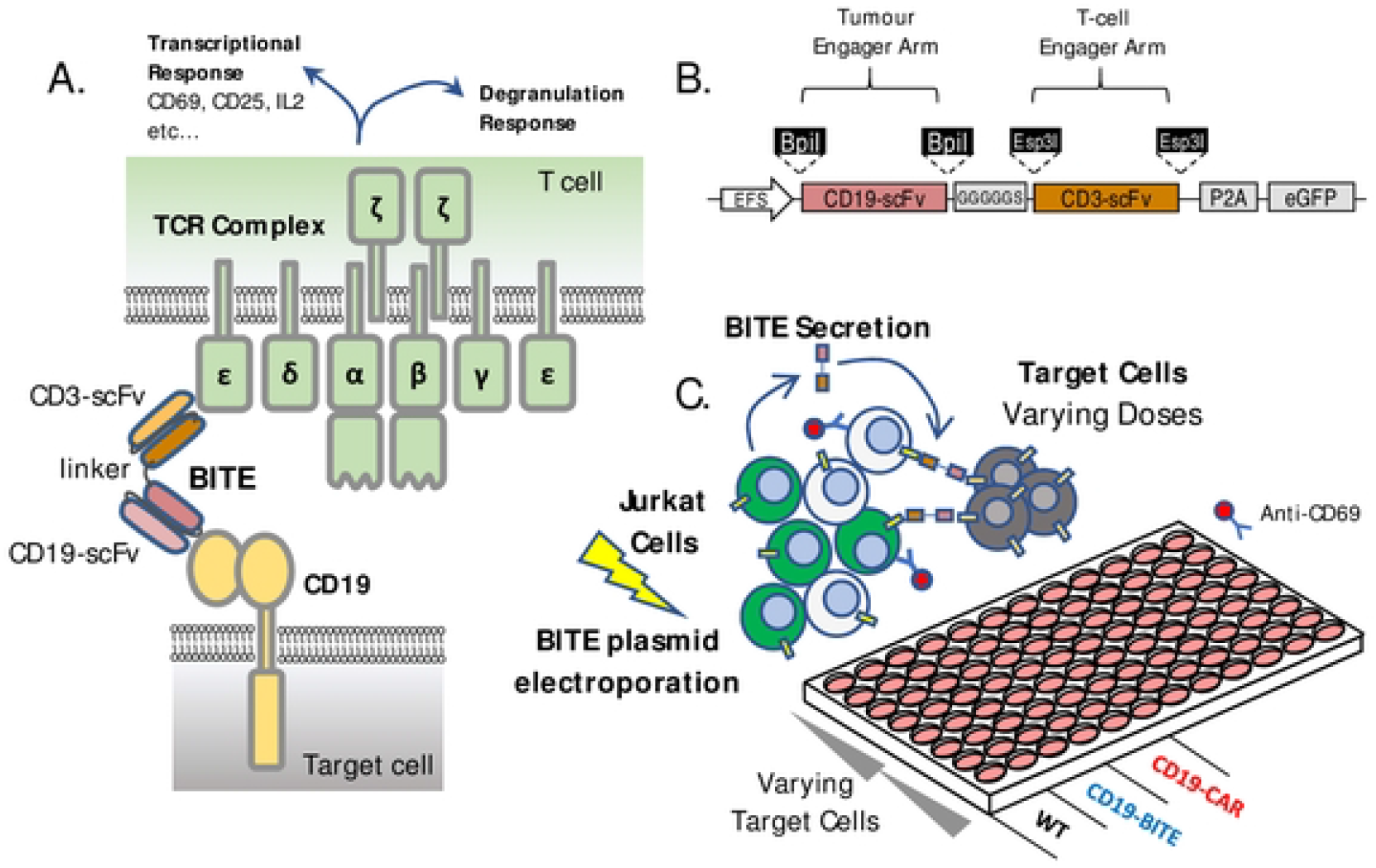
Modular BITE Plasmid and screening platform design. (A) BITE mechanism of action relies on simultaneous engagement of target cell receptors usually located on tumour cells and the CD3 domains of the T-cell receptor on T cells. This results in forming an active immune synapse between the T cell and the cancer, leading to both transcriptional and direct degranulation responses within the T cells, and target cell death. (B) The transgene design for a dual-modular BITE plasmid incorporating type-IIs restriction enzyme cassettes on each scFv region to allow easy exchange of scFv domains via golden gate cloning, (C) BITE-Jurkat screening assay setup uses direct electroporation of BITE plasmids into Jurkat cells in co-culture with various target antigen positive or negative target cell lines.

### BITE-J Function First Screening platform

Using the pBITE plasmid, we next tested a functional screening method using Jurkat electroporation to induce transient expression and secretion of the BITE molecule, similar to a method we have previously reported for screening novel chimeric antigen receptors^17^. Following BITE plasmid electroporation, Jurkat cells were co-incubated with target cells at varying effector to target ratios to produce a standard dose response curve, with Jurkat cells acting as both the BITE producer and immune effector cell (Figure 1c). After co-incubation of Jurkat and target cells, cultures were stained with anti-CD69 and analyzed via flow cytometry to assess BITE activity. While we were able to detect consistent increases in Jurkat activation with BITE expression via this method (Figure 2a), activation of Jurkat cells expressing BITE was much lower than that observed for Jurkat cells expressing a CD19-targeted CAR (Figure 2b). We hypothesized that the low activity of the BITE in this assay format might be due to relatively low electroporation efficiency in Jurkat cells, thus we wished to examine a more traditional transient production of BITE proteins in HEK293T cells.

**Figure 2:**
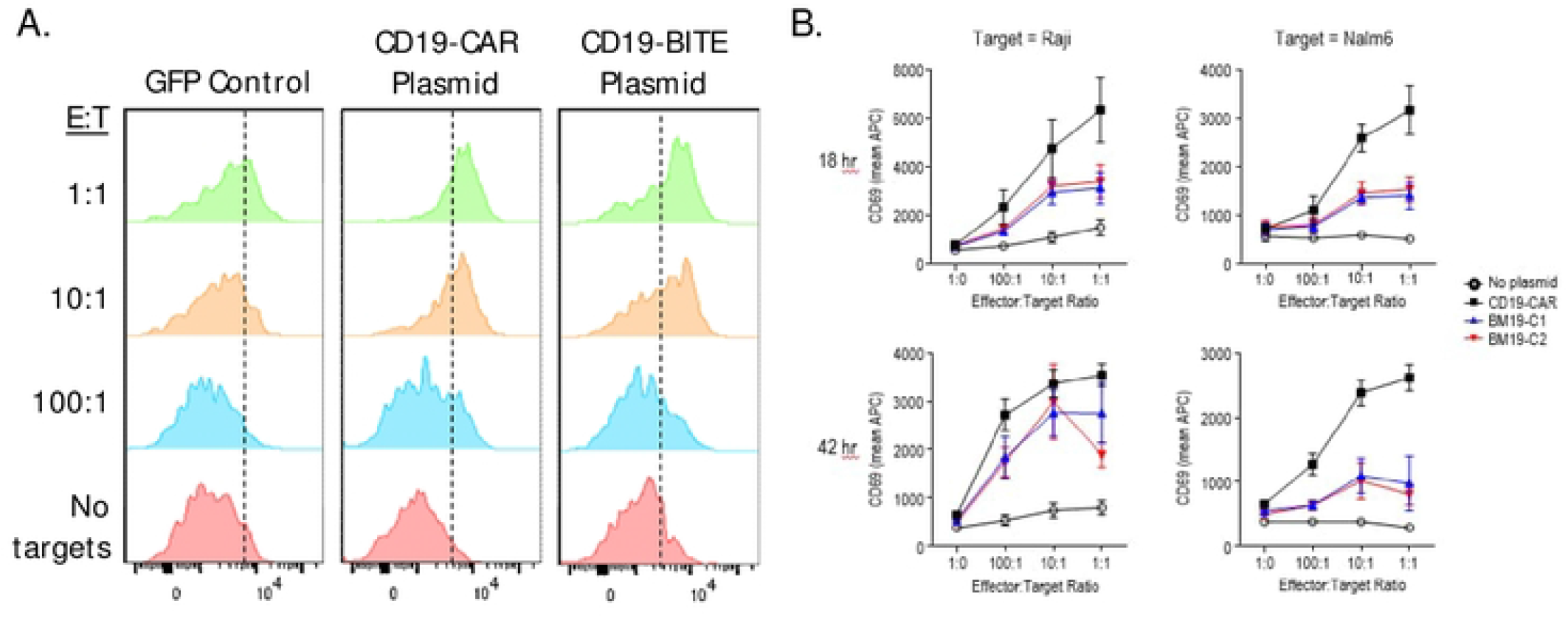
BITE-Jurkat Assay using direct electroporation in Jurkats. (A) Human Jurkat T cells were electroporated with CD19-targeted BITE or CAR plasmids as described in the methods section. Following recovery, electroporated Jurkat cells were then placed in co-culture with fluorescently labelled CD19-expressing target cells at various effector to target ratios and incubated at 37°C overnight. Co-cultures were stained with anti-human CD69-APC and analyzed via flow cytometry. Results are representative of 3 repeated experiments. (B) The mean fluorescent intensity for CD69-APC staining on gated Jurkat cells is shown for Jurkat-CD19-BITE or Jurkat-CD19-CAR cells in co-culture with CD19+ Raji or NALM6 cells for 18 hours (top) or 42 hours (bottom). Graphs show the mean result from 3 experiments repeated in duplicate +/− standard error of the mean (SEM).

### Indirect BITE-J Function First Screening platform

HEK293T cells are widely used for production of recombinant proteins where a human cell source is desirable^18^. HEK293T cells were transfected with BITE plasmids using standard PEI transfection, using fluorescence microscopy to confirm successful transfection at day 1. HEK293T cells were then grown for 5-7 days before collecting the supernatant, using filtration to clear any remaining cells. Non-purified BITE supernatants were then assayed using co-cultures of Jurkat T cells and target cells similarly to the previous iteration of the BITE-J assay (Figure 3a). These co-cultures were left for either 16 or 40 hours, then analyzed via flow cytometry for CD69 upregulation on Jurkat cells. With this improved BITE production technique, we observed more consistent BITE-induced upregulation of CD69 in the presence of increasing numbers of CD19+ Raji or NALM6 target cells (Figure 3b). The addition of non-transfected HEK293T supernatant as a control had no effect on Jurkat activation.

**Figure 3:**
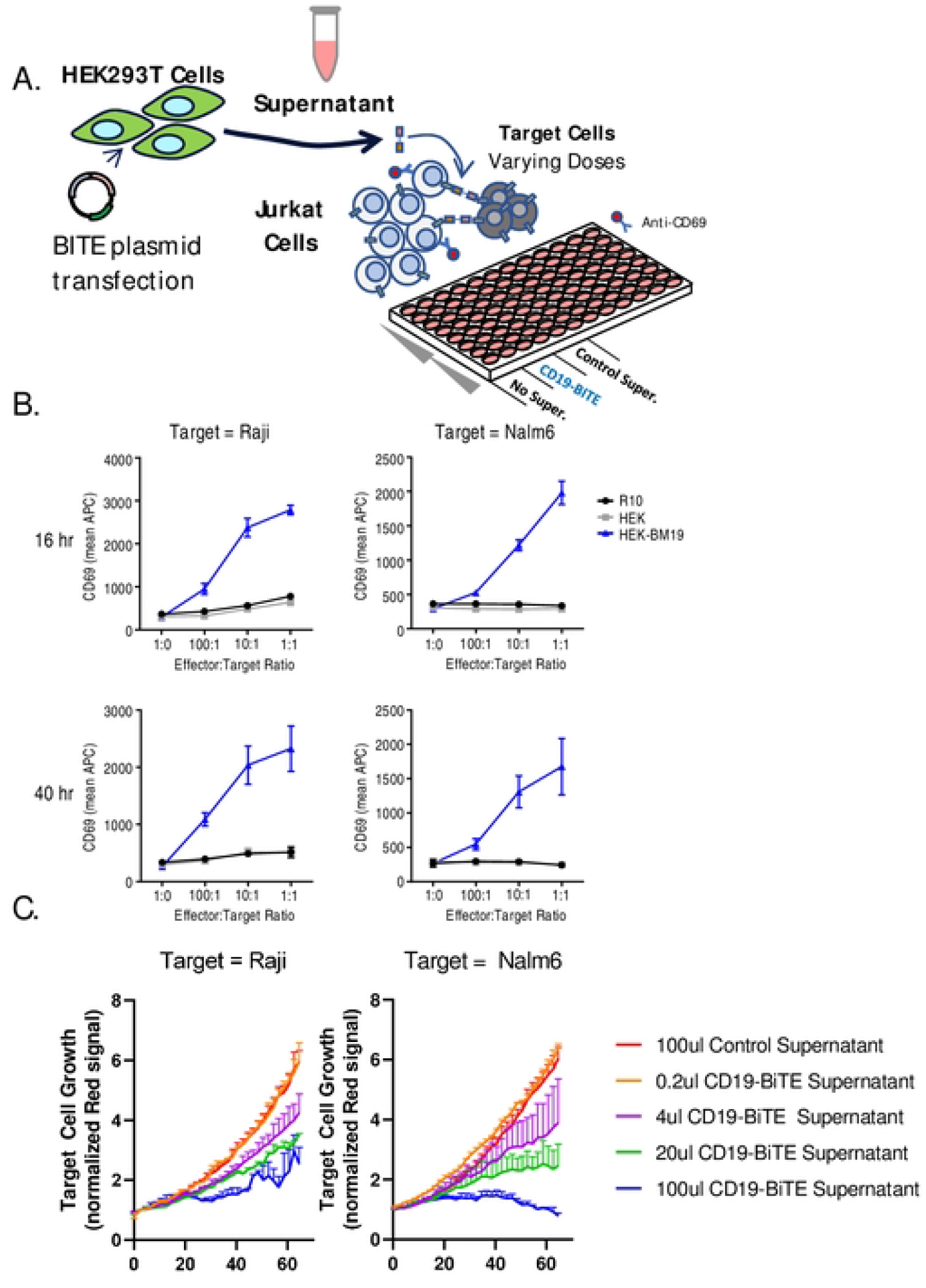
BITE-Jurkat Assay using HEK293T BITE production. (A) A schematic illustration of the workflow for production of BITE supernatant using HEK293T cells and testing using Jurkat-target cell co-cultures is shown (B) HEK293T cells were transiently transfected with BITE plasmid as described in the methods section and supernatants were collected and frozen at −80°C. 50 000 Jurkat cells were placed in a 96-well plate in co-culture with varying numbers of CD19+ Raji or NALM6 with effector to target ratios as shown. 50ul of BITE supernatant or control supernatant was then added and co-cultures were incubated at 37°C overnight. Co-cultures were stained with anti-human CD69-APC and analyzed via flow cytometry. Graphs show the mean result from 4 experiments repeated in duplicate +/− SEM. (C) Primary human T cells were placed in co-culture with 10 000 human T cells and equal numbers of red fluorescently labelled CD19+ Raji or NALM6 cells, and varying doses of CD19-BITE supernatant was added. Co-cultures were then monitored at regular intervals using fluorescence microscopy and automated cell counting via Incucyte. Graphs show the growth rate of red fluorescent target cells with varying BITE dose. Primary T cell results are from a single experiment performed in duplicate.

As this production technique yielded a large volume of BITE supernatant, much more than necessary for Jurkat experiments, we were also able to rapidly proceed to testing the effects of CD19-BITE on co-cultures of primary human T cells and CD19+ Raji or Ramos target cells. Using live fluorescence microscopy (Incucyte), we monitored the relative growth of mKate2-labelled target cells in the presence of varying doses of CD19-BITE supernatant. We observed strong dose-dependent target cell killing with BITE treatment for both target cell types (Figure 3c). These results demonstrate that transient production in HEK293T cells followed by BITE-J assay represents a viable platform for testing of biological activity for BITE molecules.

### Swapping the engager antigen

Next, we wished to test the flexibility of our BITE functional screening approach for different tumour antigen targeting domains. Using several mouse-derived single chain variable fragments (scFvs) targeting EGFRvIII, which we have previously reported to have good activity against EGFRvIII+ glioblastoma cells in CAR format^17^, we generated novel constructs using single pot restriction-ligation cloning with our BITE plasmid (Figure 4a). We then transiently transfected HEK293T cells with EGFRvIII-BITE plasmids and screened supernatants via Jurkat co-culture with EGFRvIII-positive U87vIII and DKMG glioblastoma target cell lines. Only two of the BITE plasmids generated (F269 and F263) resulted in positive CD69 upregulation when BITE supernatants were added to Jurkat/target cell co-cultures (Figure 4b). We then proceeded to testing using polyclonally expanded primary human T cells in co-culture with target antigen-positive U87-vIII target cells, or target antigen-negative U87-WT or Raji target cells. In this case, we observed rapid killing of EGFRvIII+ target cells (Figure 4c) but not EGFRvIII-negative cells (Figure 4d, e). Overall, these results demonstrate that our modular plasmid platform allows rapid cloning and screening of novel BITE molecules for T-cell redirecting activity, identifying two candidate BITE molecules F263-OKT3 and F269-OKT3 for potential future development.

**Figure 4:**
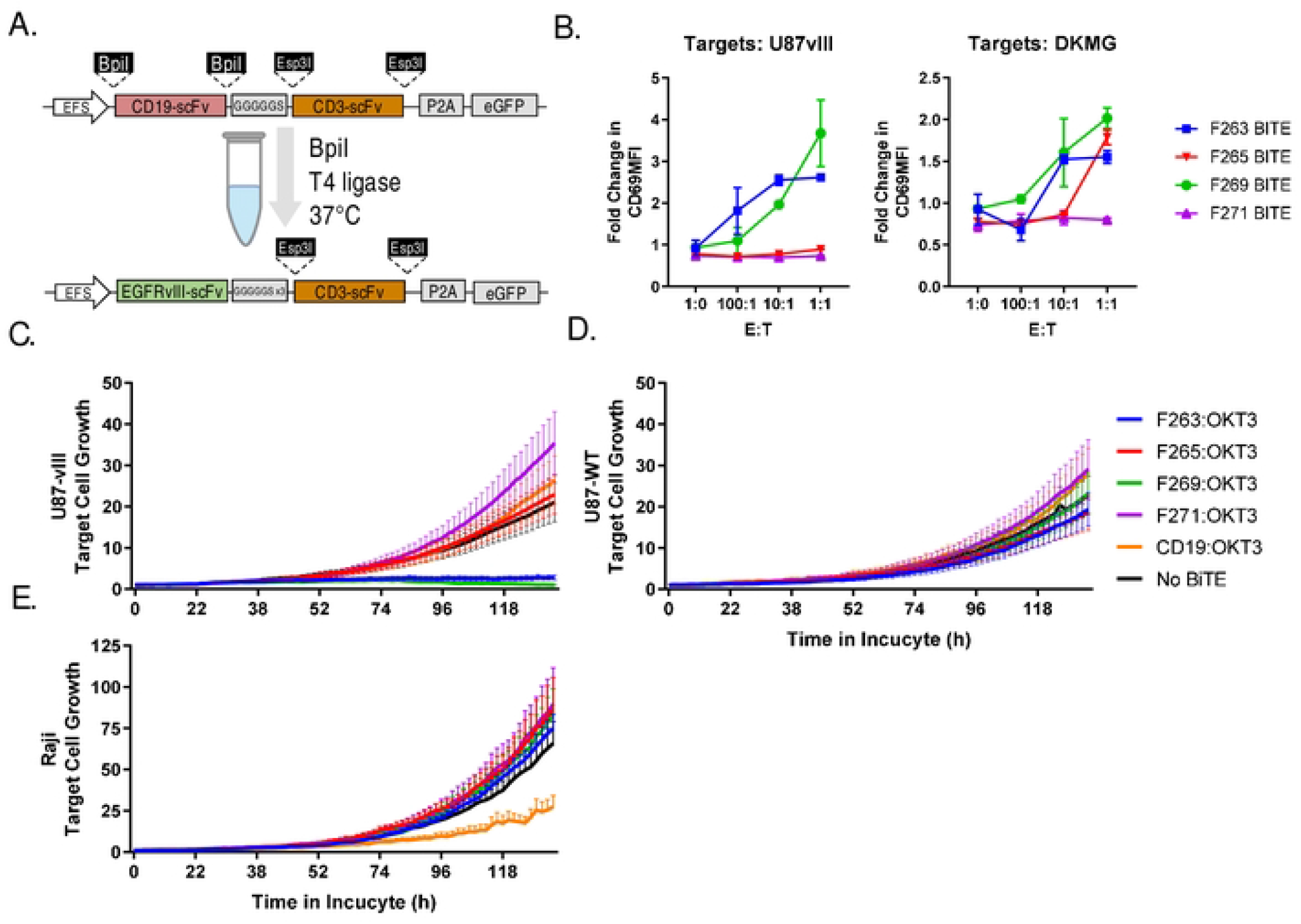
Production and screening of EGFRvIII targeted BITEs. (A) A single-pot restriction ligation reaction with BpiI restriction enzyme was used to swap CD19-targeting arm with EGFRvIII-specific scFv sequences to generate EGFRvIII-BITE plasmids as described in the methods section. (B) HEK293T cells were transiently transfected with BITE plasmids and supernatant was collected and frozen. 50 000 Jurkat cells were placed in a 96-well plate in co-culture with varying numbers of EGFRvIII+ U87vIII or DKMG target cells as shown. 50ul of BITE supernatant or control supernatant was then added and co-cultures were incubated at 37°C overnight. Co-cultures were stained with anti-human CD69-APC and analyzed via flow cytometry. Graphs show the mean result from a single experiment performed in duplicate +/− SEM. Fresh EGFRvIII or CD19-specific BITE supernatants were then generated in HEK293T cells for testing in primary T cells. 10 000 human T cells were combined with 2000 (C) EGFRvIII+ CD19-U87vIII target cells, (D) EGFRvIII-CD19-U87WT cells, or (E) EGFRvIII-CD19+ Raji cells. Graphs show the relative fluorescent signal of red-fluorescent target cells over 5 days in co-culture. Results are derived from a single experiment, but are representative of at least 3 repeated observations.

### Swapping the CD3 targeting domain

We next sought to test our platform for flexibility with respect to screening novel T-cell engaging domains of the bispecific molecule. We developed a number of novel mouse monoclonal antibodies (mAbs) against human CD3 complex using a multi-antigen immunization strategy and traditional hybridoma screening. Through this process, we were able to identify several monoclonal antibodies with reactivity to Jurkat cells (Figure 5a) and human T cells (Figure 5b). To assess whether scFvs derived from these novel CD3-targeted mAb would be functional within as part of a BITE molecule, we cloned 4 anti-human CD3 single chain variable fragments into CD19 or EGFRvIII specific BITE plasmids (Figure 5c). We then generated supernatants using transient transfection of HEK293T as described above. BITE supernatants were screened for activity using Jurkat cells in co-culture with EGFRvIII-expressing U87-VIII targets or CD19-expressing Raji cells. Previously tested constructs using OKT3 CD3-engager arms showed activity with both EGFRvIII and CD19 specific BITEs, whereas we detected activity for only one of our novel CD3-engager BITEs and only when combined with an EGFRvIII-specific scFv (Figure 5d). To confirm these results, we repeated BITE production and Jurkat co-culture screening of CD19 and EGFRvIII targeted BITE molecules incorporating OKT3 or the novel 1E2 CD3-targeting single chain variable fragment. Whereas EGFRvIII BITEs incorporating an scFv derived from the 1E2 mAb or OKT3 showed specific reactivity against EGFRvIII expressing U87-vIII cells, only CD19-OKT3 showed reactivity to CD19-expressing Raji cells (Figure 5e). These results indicate that the 1E2-scFv is functional only for an EGFRvIII-targeting BITE but not a CD19-targeting BITE.

**Figure 5:**
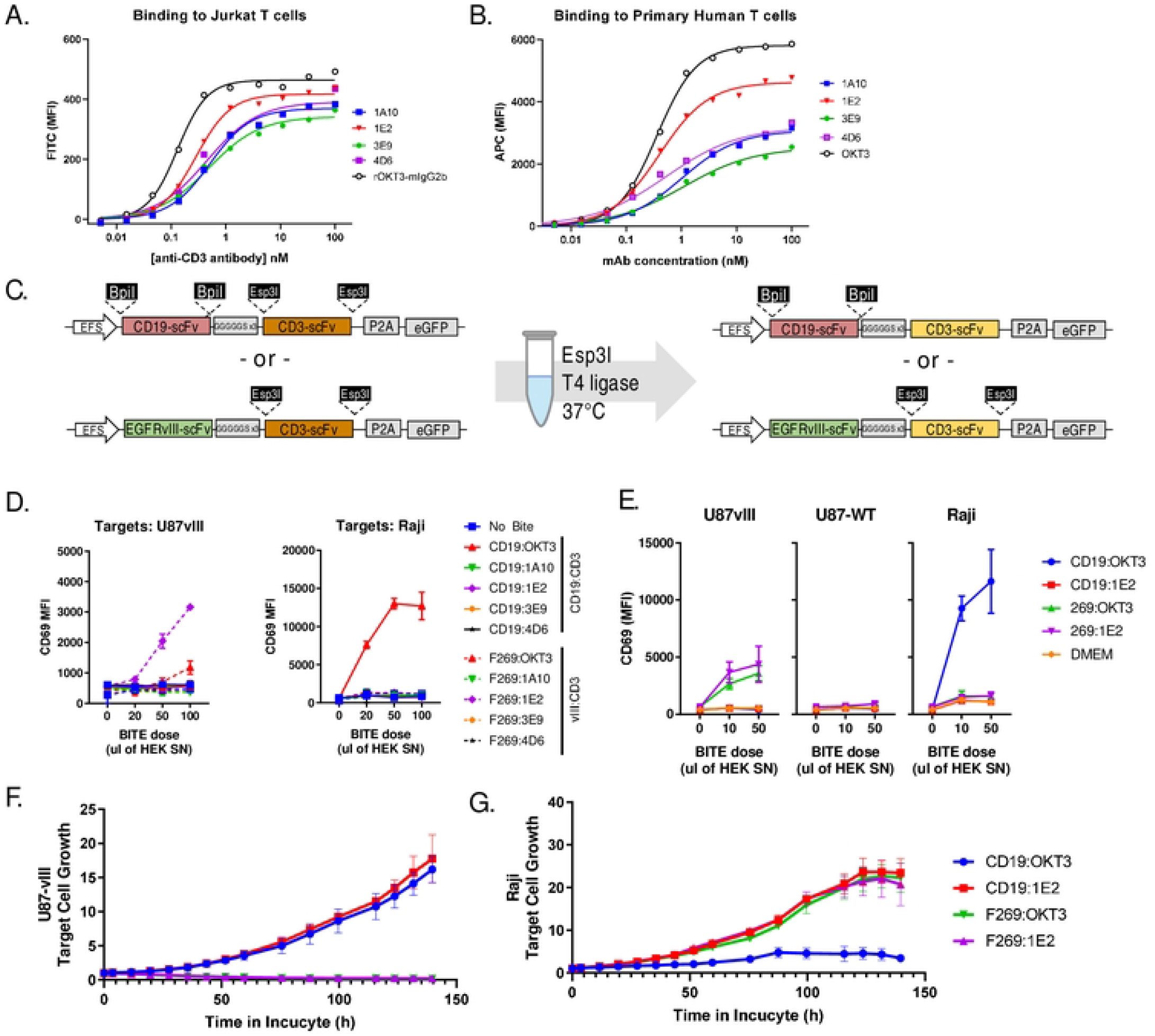
Production and screening of BITE molecules incorporating novel CD3 targeting scFvs. Novel monoclonal antibodies were generated via mouse immunization and hybridoma screening. Four monoclonal antibodies are shown which have high reactivity to (A) Jurkat T cells and (B) primary human T cells. Following this, antibody heavy and light chains were sequenced. (C) scFv DNA was then synthesized and assembled into CD19 or EGFRvIII BITE molecules using single-pot restriction ligation with Esp3I restriction enzyme. (D) Ten unique CD19 or EGFRvIII BITE plasmids were then transfected into HEK293T cells and supernatants were tested using Jurkat cells in 1:1 co-cultures with EGFR-vIII+ U87vIII cells (*left*) or CD19+ Raji cells (*right*) using varying doses of BITE supernatant as shown. (E) Four BITE plasmids found to produce active BITE molecules were tested again against EGFRvIII+ U87-vIII cells, EGFRvIII-U87-WT cells, or CD19+ Raji cells. (F) BITE supernatants for CD19 or EGFRvIII targeted molecules incorporating the 1E2 CD3-scFv were added to co-cultures with 10 000 primary human T cells and 2000 target cells. Graphs show the relative fluorescent signal of red-fluorescent target cells over 5 days in co-culture. Results are derived from a single experiment, but are representative of at least 3 repeated observations.

Finally, to assess whether this wholly novel BITE molecule can induce functional interaction between human T cells and target cells, we proceeded to screening these molecules for activity in co-cultures of polyclonally expanded human T cells with U87-VIII or Raji target cells. Human T-cells quickly killed EGFRvIII+ target cells when treated with a molecule combining EGFRvIII-scFv with OKT3 or our novel CD3-specific 1E2 scFv, but not CD19-targeted BITEs (Figure 5f, supplemental videos 1-4). In contrast, a CD19-specific scFv in combination with only OKT3 showed activity against Raji cells, but not a similar CD19-1E2 or EGFRvIII-specific BITEs (Figure 5g, supplemental videos 5-8). Overall, these results indicate that we have established a flexible platform for high-throughput functional screening of novel BITE molecules wherein either the tumour targeting or immune targeting arm can be recombined and screened for biological activity.

## Discussion

Monoclonal antibody therapies targeting many cancer-associated antigens can effectively induce antibody dependent cellular cytotoxicity (ADCC) through simultaneous engagement of target antigens via their variable domains and immune cells via the Fc domain. Many antibodies have been shown to mediate ADCC both in pre-clinical and clinical studies including CD20-targeting rituximab^19^, Her2-targeting trastuzumab^20^, or CD38 targeting daratumumab^21^. While these therapies have had remarkable success in treatment of many cancers, NK-mediated ADCC lacks the aggressively proliferative, tumour-penetrating, and inflammatory responses that can be mediated by antigen-specific T cell responses. Thus, bi-specific T cell engaging antibodies were envisioned as a means of redirecting potent T cell responses against tumours, leading to the development of blinatumomab, a CD19-CD3 BITE used to treat acute lymphoblastic leukemia, as well as other BITE therapeutics at various stages of development. While BITE therapeutics show strong potential for treatment of many different cancer types, their complex structure makes discovery and development costly and unpredictable. Thus, we sought to establish a flexible and cost-efficient platform for rapid identification of promising BITE molecules.

The system presented here, including the dual-modular BITE plasmid, transient production in HEK293T cells, and BITE-J screening of supernatants, provides a complete method for high throughput functional screening of novel BITE molecules. Other systems have been previously reported for BITE screening, but all are dependent on production of purified high-quality proteins; such as the dock and lock system used by Rossi et al. to screen multi-functional antibodies^9^, the work of Zappala et al. where antibodies are covalently linked to a CD3-specific scFv ^22^, or that by. Hofmann et al.^23^ wherein intein mediated dimerization of antibodies is used.. Whereas these methods offer a compelling means to mix and match many different binding domains for their activity in dimerized format, all require purified antibody protein, which may increase the complexity and cost of screening. Furthermore, this method may not be predictive of activity for BITEs incorporated in a single molecule for downstream production. In contrast to this, we present here a function focused approach, wherein many antibodies can be rapidly cloned and tested using transient expression. It should be noted that while the BITE-J method does not require a purification step to screen and evaluate generated BITEs, purification will ultimately be necessary for downstream testing using *in vivo* models. Given that the plasmid we developed here does not code for an Fc region, standard protein G purification is not possible. We intend to investigate alternate versions of this platform in future, for example we have developed a version of this BITE screening plasmid containing a His-tag or Fc-tag, to allow downstream purification via IMAC column.

Sugiyama et al have demonstrated a similar workflow wherein more than 52 individual bi-specific antibody molecules were generated and screened in two heavy/light chain orientations^24^. We propose that the plasmids established here provide a more flexible cloning system, as the application of simplified single-pot cloning should allow users with limited knowledge of cloning to generate novel assemblies via golden gate cloning, while also minimizing the synthesis costs for labs employing these tools. Downstream of bi-specific antibody generation and purification, most previous reports have focused on some form of T-cell Dependent Cellular Cytotoxicity (TDCC) assay^25^, typically using a viability measurements such as MTS or Luciferase reporting systems. These assays can be effective; however, we find that assessment of CD69 on Jurkat cells to be the most easily scalable method of measuring T cell activation activity. For testing with primary T cells, we find that live microscopy provides a simple method of evaluating active tumor control and killing.

Other more advanced and higher throughput approaches to screening and analysis of Bi-specific antibodies based on single cell droplet based microfluidics fluorescence sorting have also been reported^26,27^. As such approaches combine both antibody panning and antibody activation into a singular assay, they are compatible with the screening of polyclonal libraries of bi-specific molecules. The plasmids we have developed here also incorporate elements necessary for lentiviral production, and thus would be compatible with the generation of polyclonal BITE libraries. In future, we will investigate suitable methods for downstream functional assessment such as single cell sorting and screening, or microwell entrapment of BITE producing and target cells.

While the other methods of BITE screening mentioned above are all efficient high throughput methods, many of these methods require significant investment and access to specialized equipment.. BITE-J was designed not only to allow for quick BITE discovery and optimization, but also to provide tools and protocols to academic labs, which can be performed with minimal protein production and purification costs, and with minimal DNA synthesis costsBITE. We hope that the ease of access and relatively low entrance cost of BITE-J allows more researchers to contribute to the development and optimization of bi-specific therapeutics, the development of novel synthetic biology applications for BITE molecules, and to apply these tools more broadly to biological research.

The work here also extends insight into EGFRvIII-specific antibodies which we have previously reported on for CAR activity^17^. The four EGFRvIII-specific scFv molecules tested here all showed some activity in CAR format, both in the activation of CAR-expressing Jurkat cells or for inducing primary CAR-T killing of U87-VIII target cells. In contrast to this, we find that only two molecules showed significant activity here when tested in BITE format. As the screening approach employed here does not incorporate any characterization of BITE production, it is possible that low biological activity for these molecules may have been due to low productivity within HEK293T cells, and thus we cannot state conclusively that these molecules would not be functional if produced and purified in other formats.

Similarly, we also find that only one of the novel CD3-targeting scFvs tested here showed activity in BITE format, and even then, activity was restricted to combination with EGFRvIII-targeting scFv and not with a CD19-specific scFv. Again, we have no insight into the failure rate for BITE molecules in this assay, as the intent of this assay is to provide a rapid means for testing biological activity rather than focusing on various aspects of antibody characyerization. We are currently undertaking molecular studies to test whether incorporating different linker domains may be able to improve the activity of BITEs using the novel CD3-scFv reported to have activity here, something that may provide further insight into the design constraints for these BITE molecules.

The fully novel EGFRvIII-CD3 engaging molecule identified at the end of the manuscript provides a proof of principle example for high throughput discovery of novel BITE therapeutics. As this molecule shows strong activity in mediating T-cell killing of antigen-positive tumour cells, it also shows potential for downstream development. In the future, we intend to expand on this platform to better understand the design space and study the biology of bi-specific antibody therapeutics that mediate active engagement of T-cells or other immune cells with target cells.

## References

1. Weiner, G. J. Monoclonal antibody mechanisms of action in cancer. Immunol. Res. 39, 271–278 (2007).

2. Tian, Z., Liu, M., Zhang, Y. & Wang, X. Bispecific T cell engagers: an emerging therapy for management of hematologic malignancies. J. Hematol. Oncol.J Hematol Oncol 14, 75 (2021).

3. Study of the BITE® Blinatumomab (MT103) in Adult Patients With Relapsed/Refractory B-Precursor Acute Lymphoblastic Leukemia (ALL) - Study Results - ClinicalTrials.gov. https://clinicaltrials.gov/ct2/show/results/NCT01209286.

4. Wu, J., Fu, J., Zhang, M. & Liu, D. Blinatumomab: a bispecific T cell engager (BITE) antibody against CD19/CD3 for refractory acute lymphoid leukemia. J. Hematol. Oncol.J Hematol Oncol 8, 104 (2015).

5. Huehls, A. M., Coupet, T. A. & Sentman, C. L. Bispecific T cell engagers for cancer immunotherapy. Immunol. Cell Biol. 93, 290–296 (2015).

6. Wang, Q. et al. Design and Production of Bispecific Antibodies. Antibodies Basel Switz. 8, E43 (2019).

7. Mechanism of Action | BLINCYTO® (blinatumomab). https://www.blincytohcp.com/mrd/moa.

8. Viardot, A., Locatelli, F., Stieglmaier, J., Zaman, F. & Jabbour, E. Concepts in immuno-oncology: tackling B cell malignancies with CD19-directed bispecific T cell engager therapies. Ann. Hematol. 99, 2215–2229 (2020).

9. Rossi, E. A. et al. Stably tethered multifunctional structures of defined composition made by the dock and lock method for use in cancer targeting. Proc. Natl. Acad. Sci. U. S. A. 103, 6841–6846 (2006).

10. Blinatumomab. https://go.drugbank.com/drugs/DB09052.

11. Gibson, D. G. et al. Enzymatic assembly of DNA molecules up to several hundred kilobases. Nat. Methods 6, 343–345 (2009).

12. Gibson Assembly - OpenWetWare. https://openwetware.org/wiki/Gibson_Assembly.

13. Mccomb, S. et al. Antigen-Binding Agents That Specifically Bind Epidermal Growth Factor Receptor Variant Iii. (2020).

14. Marcil, A., Jaramillo, M., Sulea, T., Moreno, M. & Wu, C. Anti-Egfrviii Antibodies and Antigen-Binding Fragments Thereof. (2020).

15. Molla Kazemiha, V. et al. PCR-based detection and eradication of mycoplasmal infections from various mammalian cell lines: a local experience. Cytotechnology 61, 117–124 (2009).

16. Chicaybam, L., Sodre, A. L., Curzio, B. A. & Bonamino, M. H. An Efficient Low Cost Method for Gene Transfer to T Lymphocytes. PLOS ONE 8, e60298 (2013).

17. Bloemberg, D. et al. A High-Throughput Method for Characterizing Novel Chimeric Antigen Receptors in Jurkat Cells. Mol. Ther. - Methods Clin. Dev. 16, 238–254 (2020).

18. Tan, E., Chin, C. S. H., Lim, Z. F. S. & Ng, S. K. HEK293 Cell Line as a Platform to Produce Recombinant Proteins and Viral Vectors. Front. Bioeng. Biotechnol. 9, 796991 (2021).

19. Reff, M. E. et al. Depletion of B cells in vivo by a chimeric mouse human monoclonal antibody to CD20. Blood 83, 435–445 (1994).

20. Spector, N. L. & Blackwell, K. L. Understanding the Mechanisms Behind Trastuzumab Therapy for Human Epidermal Growth Factor Receptor 2–Positive Breast Cancer. J. Clin. Oncol. 27, 5838–5847 (2009).

21. Casneuf, T. et al. Effects of daratumumab on natural killer cells and impact on clinical outcomes in relapsed or refractory multiple myeloma. Blood Adv. 1, 2105–2114 (2017).

22. Zappala, F. et al. Rapid, site-specific labeling of “off-the-shelf” and native serum autoantibodies with T cell–redirecting domains. Sci. Adv. 8, eabn4613 (2022).

23. Hofmann, T. et al. Intein mediated high throughput screening for bispecific antibodies. mAbs 12, 1731938 (2020).

24. Sugiyama, A. et al. A semi high-throughput method for screening small bispecific antibodies with high cytotoxicity. Sci. Rep. 7, 2862 (2017).

25. Nazarian, A. A. et al. Characterization of bispecific T-cell Engager (BITE) antibodies with a high-capacity T-cell dependent cellular cytotoxicity (TDCC) assay. J. Biomol. Screen. 20, 519–527 (2015).

26. Wang, Y. et al. High-throughput functional screening for next-generation cancer immunotherapy using droplet-based microfluidics. Sci. Adv. 7, eabe3839 (2021).

27. Segaliny, A. I. et al. A high throughput bispecific antibody discovery pipeline. 2021.09.07.459213 (2021) doi:10.1101/2021.09.07.459213.

